# Resolving genotype-phenotype discrepancies of the Kidd blood group system using long-read nanopore sequencing

**DOI:** 10.1101/2023.05.13.540649

**Authors:** M Gueuning, GA Thun, N Trost, L Schneider, S Sigurdardottir, C Engström, N Larbes, Y Merki, BM Frey, C Gassner, S Meyer, MP Mattle-Greminger

## Abstract

Due to substantial improvement in read accuracy, third-generation long-read sequencing holds great potential in blood group diagnostics, particularly in cases where traditional genotyping or sequencing techniques, primarily targeting exons, fail to explain serological phenotypes. In this study, we employed Oxford Nanopore sequencing to resolve all genotype-phenotype discrepancies in the Kidd blood group system (JK, encoded by *SLC14A1*) observed over seven years of routine high-throughput donor genotyping using a mass spectrometry based platform at Blood Transfusion Service Zurich. Discrepant results from standard serological typing and donor genotyping were confirmed by commercial PCR-SSP kits. To resolve discrepancies, we amplified the entire coding region of *SLC14A1* (∼24 kb, exons 3 to 10) in two overlapping long-range PCRs in all samples. Amplicons were barcoded and sequenced on a MinION flow cell. Sanger sequencing and bridge-PCRs were used to confirm findings. Among 11,972 donors with both serological and genotype data available for the Kidd system, we identified 10 cases with unexplained conflicting results. Five were linked to known weak and null alleles caused by variants not included in the routine donor genotyping. In two cases, we identified novel null alleles on the *JK*01* (Gly40Asp; c.119G>A) and *JK*02* (Gly242Glu; c.725G>A) haplotype, respectively. Remarkably, the remaining three cases were associated with a yet unknown deletion of ∼5 kb spanning exons 9-10 of the *JK*01* allele, which other molecular methods had failed to detect. Overall, nanopore sequencing demonstrated reliable and accurate performance for detecting both single-nucleotide and structural variants. It possesses the potential to become a robust tool in the molecular diagnostic portfolio, particularly for addressing challenging structural variants such as hybrid genes, deletions and duplications.

## Introduction

The latest advancements in third-generation long-read sequencing (TGS) technologies offer notable advantages, including the capacity to elucidate extensive haplotypes and characterize genomic regions that pose challenges to Sanger sequencing or next-generation short-read sequencing [1]. In the specific field of transfusion medicine, TGS complements traditional approaches used for the blood group assessment of samples with complex or discordant serological and genetic results [2–6]. Such cases include serological reactions exhibiting unexpected weak agglutination or null phenotypes, often resulting from rare or unknown genetic variants that are not typed in routine genotyping workflows [7].

Traditionally, Sanger sequencing of the underlying blood group genes has been used to resolve such elusive cases. However, it has several limitations. A major drawback is the lack of phase information, i.e. the inability to reconstruct haplotypes, due to overlapping signals from maternal and paternal alleles. This makes, for instance, functional interpretation challenging since identified potentially causative variants cannot be assigned to the respective blood group allele background. Another limitation is that indels and structural variants (SVs) cause a shift in signals and thus hamper the readability of the sequences. Length restriction of Sanger sequencing is another limitation, which usually results in neglecting the investigation of non-coding regions of a gene, including the promoter region, enhancer elements, and transcription factor binding sites. The limited length of the sequenced region also hinders the detection of SVs. Large deletions, for example, may simply go unnoticed with Sanger sequencing in case of an unaffected second allele in the background, which gets sequenced (allelic dropout). Some of these limitations would be mitigated when using short-read sequencing technologies, but the short read length still hampers full-gene haplotype reconstruction and the analysis of SVs, beside the lack of scalability to single sample diagnostics.

The long-read sequencing technology by Oxford Nanopore Technologies (ONT) represents a promising solution to overcome limitations of aforementioned sequencing approaches. It allows sequencing long reads of blood group genes at the kilobase (kb) scale in a low throughput setting at high quality [8]. This greatly facilitates haplotype reconstruction and the inclusion of introns and proximal regulatory elements [5,9]. Moreover, it lowers the risk of misinterpreting loss of heterozygosity caused by allelic dropout as the long reads are much more likely to cover the respective breakpoint sites. Previous constraints of ONT sequencing, in particular the low single-read accuracy and the need for advanced bioinformatics skills [10–12], have been greatly alleviated by latest developments.

In this study, we propose a cost-effective and reliable approach based on ONT sequencing to resolve the Kidd blood group (JK) in samples with discordant serological and genotypic findings. We used the JK system as a proof-of-concept due to a combination of its clinical relevance and its low antigen diversity, enabling straightforward phenotyping. In fact, this blood group system only comprises three antigens: the antithetical JK1 and JK2 antigens, also known as Jk(a) and Jk(b), as well as the high-prevalence antigen JK3 [13]. Antibody formation by alloimmunization is particularly common in patients with sickle cell disease [14] and JK antibodies are a cause of delayed hemolytic transfusion reactions as well as hemolytic disease of the newborn [15]. The *SLC14A1* gene, spanning over ∼30 kb on chromosome 18, consists of 10 exons and translates into a 389 amino acid glycoprotein. The International Society of Blood Transfusion (ISBT) currently recognizes over 60 different *JK* alleles, of which most represent null alleles caused by missense or nonsense single-nucleotide variants (SNVs) [13].

Here, we investigated all unresolved genotype-phenotype discrepant cases collected over seven years of routine high-throughput donor genotyping at Blood Transfusion Service Zurich (Switzerland) using Matrix-Assisted Laser Desorption Ionization – Time-of-Flight mass spectrometry (MALDI-TOF MS) [7,16]. The workflow was based on long-range PCR (LR-PCR) amplification of *SLC12A1* and subsequent barcoded nanopore-sequencing of amplicons. As only the LR-PCR primer design was gene-specific, our workflow can easily be customized to target other blood group genes. Overall, our results demonstrated that ONT sequencing can effectively and accurately resolve genotype-phenotype discrepancies, also those unresolved by Sanger sequencing. The cost-effective approach holds great promise in general for accurate blood group determination in challenging cases.

## Material and methods

A graphical overview of how discrepant samples were identified and processed is provided in Figure 1.

**Figure 1.**
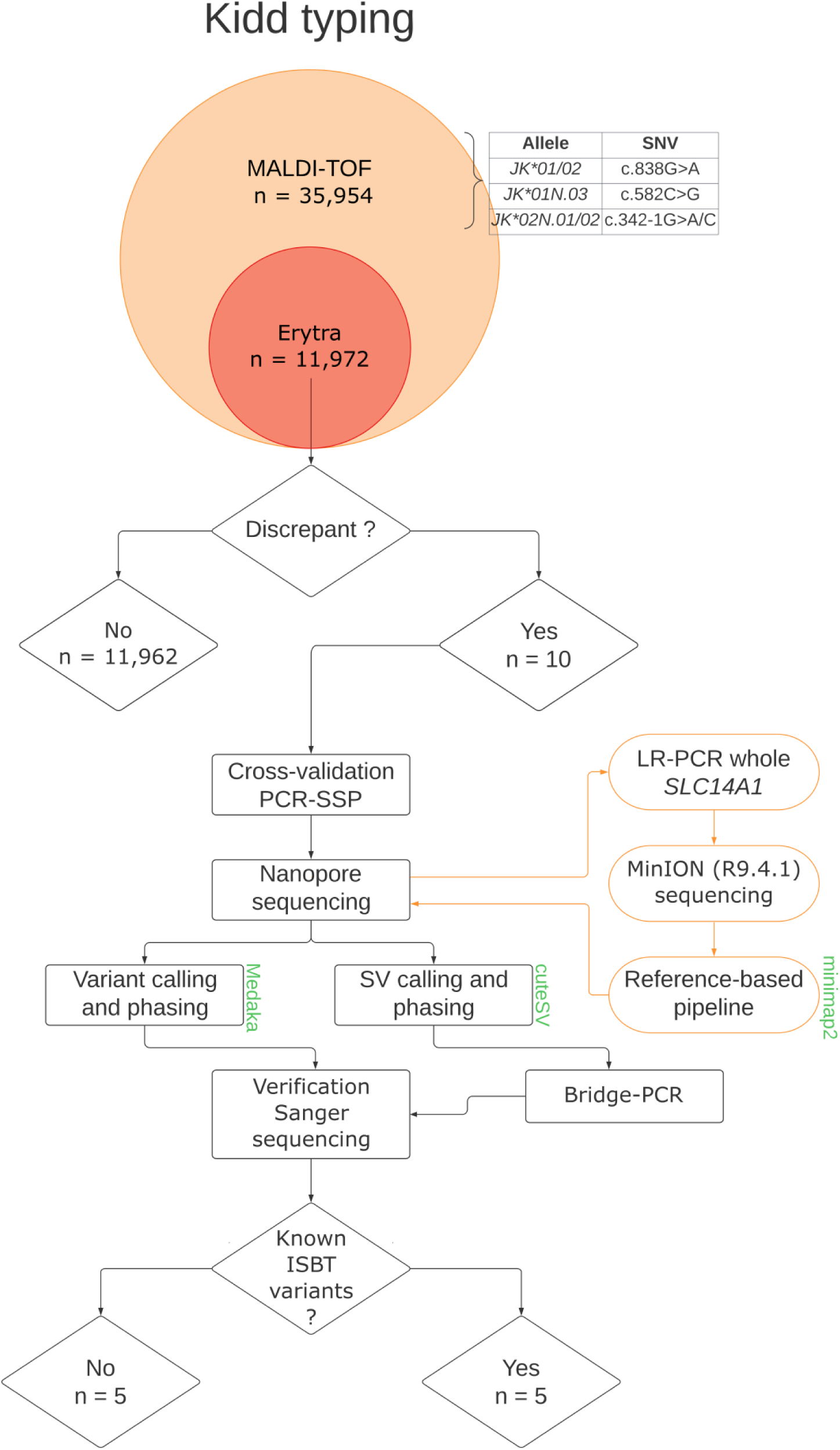
Flowchart depicting identification and processing of genotype-phenotype discrepancies. Main bioinformatics tools are provided in green.

### Routine high-throughput donor geno- and phenotyping

Genomic DNA of blood donors was extracted on a Chemagic MSM I extraction robot (Chemagen, Perkin Elmer). High-throughput genotyping of 46 blood group antigens of 35,954 donor samples collected in the years 2015 to 2021 was performed by MALDI-TOF MS as described previously [16,17]. Genotyping included *JK* alleles of interest in a Swiss and international donor pool. Specifically, *JK*01* and *JK*02* were discriminated by the SNV c.838G>A, the null alleles *JK*01N.03* (found in Swiss families) [18], *JK*02N.01* (predominantly detected in Asian populations [19]), and *JK*02N.02* were typed using the c.582C>G, c.342-1G>A, and c.342-1G>C variants [20], respectively. All coding DNA positions provided are based on reference transcript NM_015865.7.

Of all donors genotyped by MALDI-TOF MS, 11,972 donors were additionally serologically phenotyped on a follow-up donation using Erytra automated system (Grifols) for seven blood groups, among which was JK. In case of genotype-phenotype discrepancy between MALDI-TOF MS genotyping and phenotyping on the Erytra, the sample was analyzed further using other standard serological techniques and commercial PCR-SSP kits (sequence-specific-priming PCR; inno-train). For this, DNA was manually re-extracted from the follow-up donation using the Nucleon BACC 3 kit (Gen-Probe Life Sciences). DNA concentrations were measured using Nanodrop 2000 (Thermo Fisher Scientific).

### Nanopore sequencing of genotype-phenotype discrepancies

In case of confirmed genotype-phenotype discrepancy, the entire coding region of the *SLC14A1* gene (∼24 kb, exon 3 to 10) was amplified and sequenced using ONT. Amplifications were done with two LR-PCRs. The first fragment (12,981 bp) was amplified using the forward primer 5’-TGCCACTTGAGTGTTTTCATTTGATGCTGC-3’ and reverse primer 5’-CACTATCCCTCCTCCTTTTTGTTCCCAAGC-3’. PCR primers for the second fragment (13,112 bp) were 5’-AAGTGACGTCCCCTCTCTGAGAGCATTAAA-3’ (forward) and 5’-AACATTCTGACAAGTGGCTGGTCCTAGAGA-3’ (reverse). To facilitate subsequent phasing, both LR-PCR fragments were designed to have a large overlap (∼1,550 bp). PCR amplifications were done in duplicates using the PrimerSTAR GXL polymerase (Takara Bio) and 250 ng of genomic DNA per reaction, following the manufacturer’s protocol. To increase amplification success, we added 1 M of Betaine enhancer (VWR) per reaction. The PCR’s profile followed a 2-step approach with a 10 second denaturation step at 95°C and a 10 minute extension step at 68°C for 30 cycles. Amplification success was verified on a 0.8% agarose gel stained with GelRed Nucleic Acid Gel (Biotium). After verification, PCR replicates were pooled and purified with 1x Agencourt AMPure XP magnetic beads (Beckman Coulter). Purified PCR products were eluted in 25 µl sterile H2O and quantified using a dsDNA broad range assay kit on a Qubit fluorometer 3.0 (both Invitrogen). Sequencing libraries were constructed following ONT’s ’Amplicon barcoding with Native Barcoding’ protocol (version: NBA_9102_v109_revF_09Jul2020). As starting material, we pooled 50 fmol of both purified amplicons for each sample. After completion of the protocol, 44 fmol of the final barcoded library was loaded on a MinION (R9.4.1) flow cell. Sequencing was stopped after no more pores were active (∼72 h).

### Nanopore bioinformatics processing

Raw nanopore reads were demultiplexed and basecalled using ONT’s *Guppy* (v.4.4.2). After demultiplexing, reads in FASTQ format were filtered based on expected length and observed read length distributions. More specifically, we filtered out all reads shorter than 12,500 bp and longer than 13,600 bp. For three samples (s02, s03 and s07) harboring a second shorter peak in read length distribution (∼8,000 bp; Figure 2c) we additionally kept all reads having a length between 7,500 bp and 8,500 bp. After size selection, *PoreChop* (v.0.2.3) was used to trim remaining adapter and barcode sequences.

**Figure 2.**
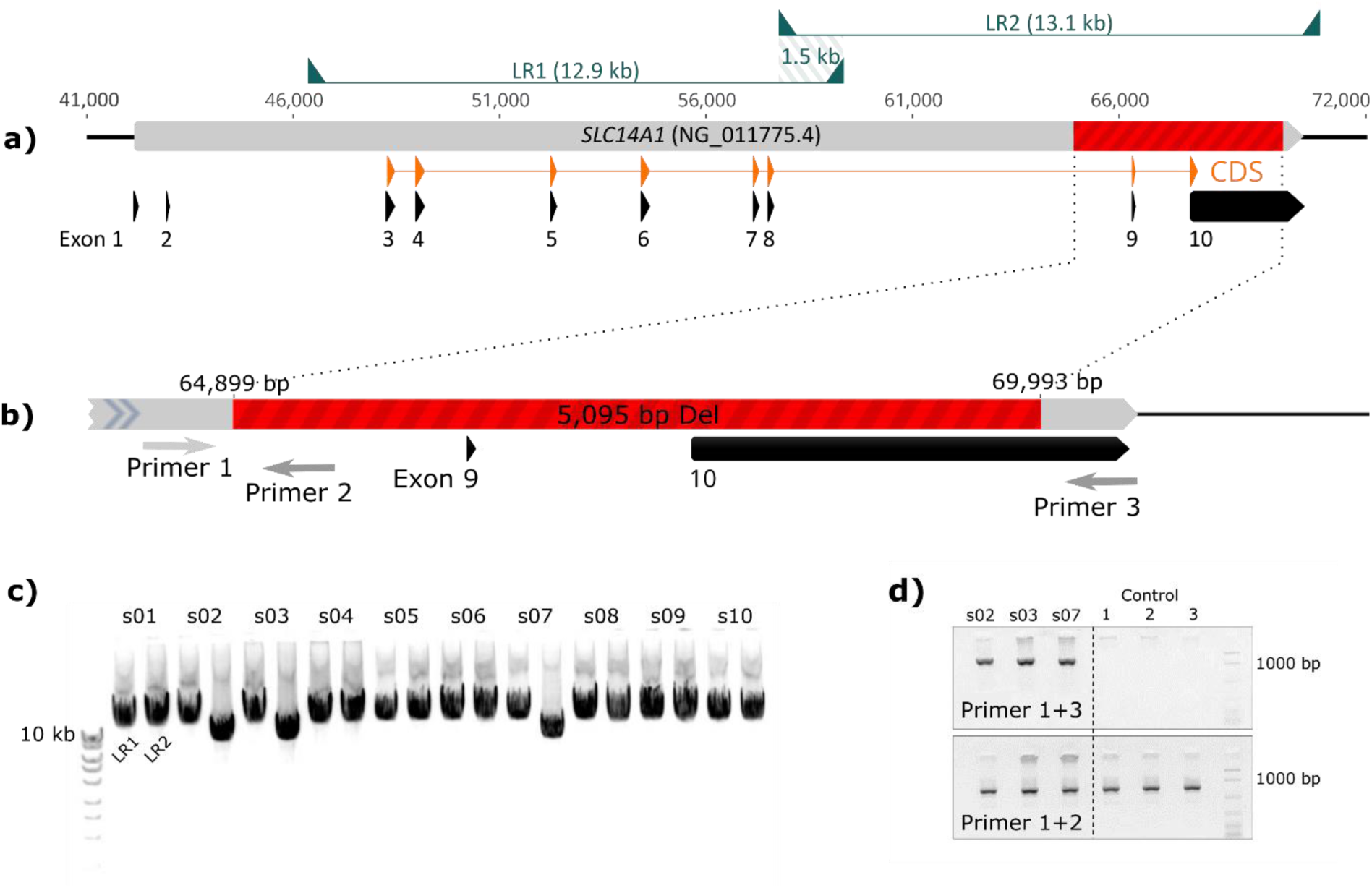
Genetic structure of the *SLC14A1* gene encoding the JK blood group system, and details on the identified large novel deletion. (a) *SLC14A1* gene showing exons, coding DNA sequence (CDS), newly identified ∼5 kb deletion (red), positions of long-range PCRs (LR1/LR2), as well as overlap between both amplicons. (b) Region of ∼5 kb deletion with breakpoints and primer positions used for bridge-PCR. (c) Gel electrophoresis of long-range PCR products (LR1 and LR2) of all samples. (d) Gel electrophoresis of bridge-PCR products for the three samples with the novel deletion and wild-type control reactions.

To reduce computational burden, reads were randomly downsampled to 1000 reads per fragment using *seqkit* (v.0.15). To differentiate between fragment 1 and 2, which were very similar in read length, all size-selected reads were first mapped to the *SLC14A1* reference sequence (NG_011775.4) using *minimap2* (v.2.17) and separated according to their mapped genomic location. For the three samples showing the secondary shorter peak, mapping showed that reads responsible for this peak partially corresponded to fragment 2. Therefore, we further downsampled to 500 reads for both the long and short fragment 2. Finally, uniformly downsampled datasets were remapped to NG_011775.4 using *minimap2*, allowing the presence of supplementary alignments (i.e. split-read mapping). Mapped reads were sorted, indexed and converted to BAM format using *samtools* (v.1.15).

Variant calling and phasing of called variants was performed using *Medaka* (v.1.2.3). As a consequence of the high coverage, threshold quality for calling indels and SNVs (default: Q9 and Q8, respectively) was increased to Q10. Presence of SVs was investigated with *cuteSV* (v.1.0.11). Command options were set as recommended for nanopore data. *BCFtools* (v.1.9) was finally used to output phased sequences in FASTA format.

### Variant confirmation by Sanger sequencing and bridge-PCR

Nanopore sequencing results were confirmed using Sanger sequencing. Specific PCR primers for each newly detected variant were designed, and PCRs were carried out using AmpliTaq DNA polymerase (Life Technologies). Primers and PCR protocols are available on request. Amplicon purification and Sanger sequencing were outsourced to an external company (Microsynth AG).

Finally, a bridge-PCR assay was designed to confirm a large deletion found with nanopore sequencing in three samples. One primer pair was designed to amplify over the deletion using primers located at the flanking regions, with the forward primer being located upstream of the 5’ breakpoint and the reverse primer downstream of the 3’ breakpoint. Another reverse primer was designed to amplify the wild type allele, enabling assessment of the heterozygosity status of the deletion. This reverse primer was designed to bind within the region of the deletion and amplify with the upstream primer targeting the 5’ breakpoint. For both primer pairs, three wild type controls were used.

## Results

### Observed genotype-phenotype discrepancies

Congruence of JK phenotypes deduced from genotypes of seven years of high-throughput donor screening and serological typing was very high. In detail, of the 35,954 donors genotyped by MALDI-TOF MS, 35,937 (99.95%) passed quality control and were assigned to *JK*01* and/or *JK*02* alleles using the SNV c.838G>A. Among them, 50.43% (n = 18,124) were *JK*01/02* heterozygous, 26.30% (n = 9,452) *JK*01/01* homozygous, and 23.27% (n = 8,361) *JK*02/02* homozygous, as expected under Hardy-Weinberg equilibrium (P>0.05). Out of 11,972 donors with available serological data, we observed seven samples where the null allele causing variants included in the MALDI-TOF MS module explained their null phenotype (n = 4 for *JK*01N.03* and n = 3 for *JK*02N.01*). There were only 10 discrepant cases (0.08%) in which MALDI-TOF MS genotyping did not agree with the observed phenotype. Phenotypes and genotypes in all of them were confirmed by additional serological analyses and PCR-SSP.

### Nanopore sequencing output

We obtained very high coverage for each amplified fragment in all 10 samples with discordant serology and genotype (Figure 3). Overall, the nanopore sequencing run produced ∼1 million reads (≥ Q7) with a median PHRED score of 12.7. After demultiplexing and filtering reads according to fragment sizes, mean number (± standard deviation) of reads per barcode (i.e. sample) was 41,640 (± 15,182).

**Figure 3.**
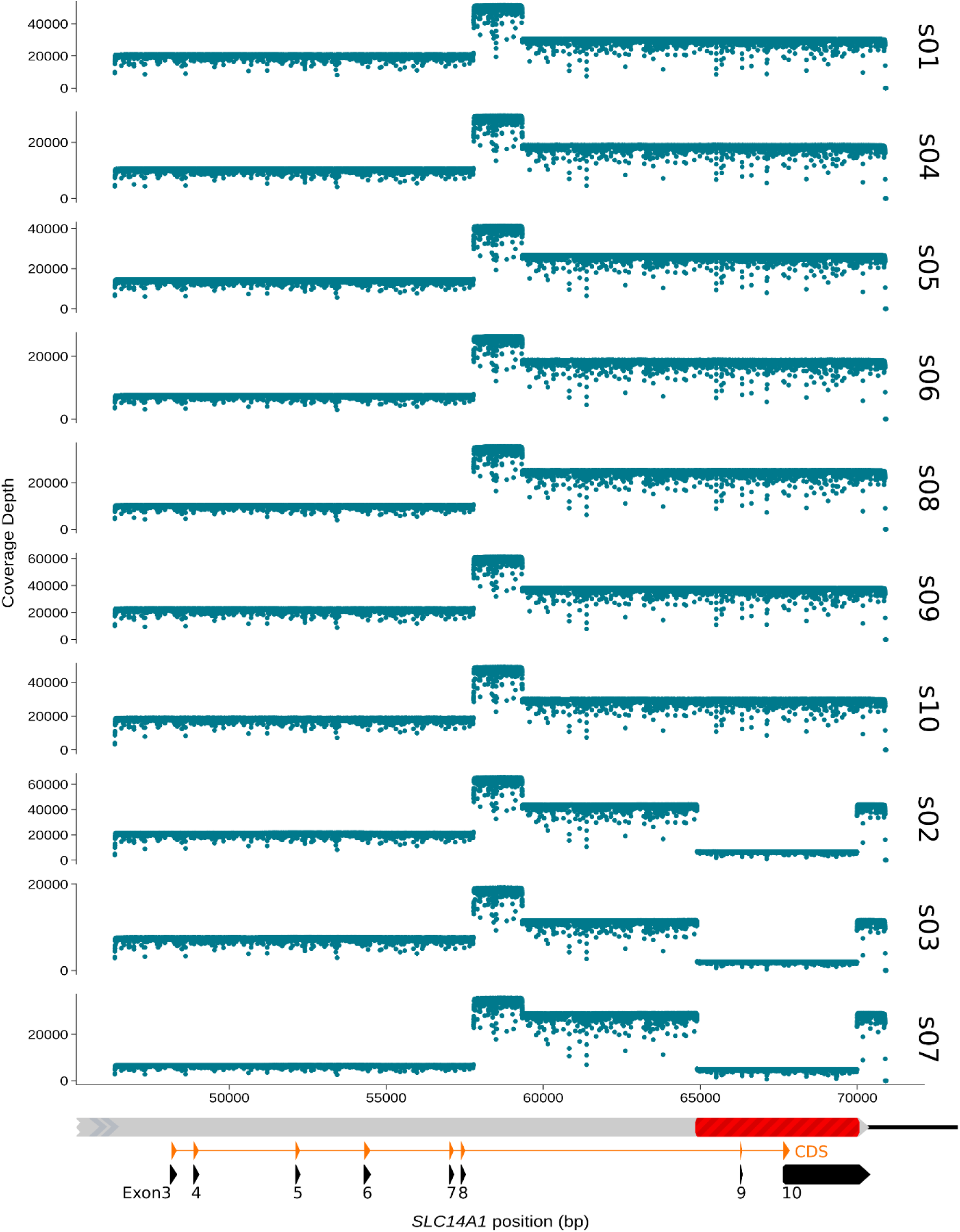
Nanopore sequencing coverage by nucleotide position for each sample. Coverages were computed by mapping length-filtered reads to *SLC14A1* reference sequence (NG_011775.4) using minimap2. For readability, samples showing a drop in coverage in LR2 were grouped together. Graphical representation of mapping location on *SLC14A1* gene is given underneath the coverage plots. Coverage scales are not fixed.

Mapping of filtered reads against the *SLC14A1* reference sequence showed a drop in coverage over a length of 5 kb in the second fragment for three samples (Figure 3). For these same samples (s02, s03 and s07), the gel electrophoresis for verifying PCR amplification successes prior to sequencing had also already revealed the presence of shorter fragments in the second LR-PCR (Figure 2c).

### Resolving discrepant cases

A summary of serology and genetic characterization of the 10 discrepant cases is provided in Table 1. In seven cases we detected SNVs in coding regions that could explain the observed phenotype. In five donors, these variants had previously been reported causing weak *(JK*01W.05*, *JK*02W.03*) and null alleles (*JK*02N.06/08/09*), respectively. In the other two donors, we identified exonic SNVs that had so far not been linked to Kidd phenotypes. In one case, the SNV was located in exon 3 (c.119G>A; NG_011775.4:g.48415G>A) on a *JK*01* allele, altering the protein sequence close to the N-terminus (Gly40Asp). The other novel SNV was located in exon 7 (c.725G>A, NG_011775.4:g.57200G>A) on a *JK*02* allele and corresponded to a missense mutation changing the codon 242 from glycine to glutamic acid (Gly242Glu). Sanger sequencing confirmed the presence of all those SNVs.

**Table 1.** Summary of serology, genotyping, and nanopore sequencing for all ten genotype-phenotype discrepancy cases. Discrepant antigens, based on comparison of observed with deduced phenotype, as well as causative alleles are highlighted in red.

| Sample | Observed serology | Deduced phenotype<br>based on MALDI-TOF MS<br>genotyping | Nanopore sequencing |  |
| --- | --- | --- | --- | --- |
|  |  |  | Haplotype 1 | Haplotype 2 |
| Known weak and null alleles |  |  |  |  |
| s01 | Jk(a+<sup>weak</sup> b-) | Jk(a+b+) | *JK\*01W.06* | *JK\*02N.08<sup>†</sup>* |
| s04 | Jk(a+b-) | Jk(a+b+) | *JK\*01* | *JK\*02W.03* |
| s05 | Jk(a-b+) | Jk(a+b+) | *JK\*01W.05* | *JK\*02<sup>†</sup>* |
| s06 | Jk(a+b-) | Jk(a+b+) | *JK\*01* | *JK\*02N.06<sup>†</sup>* |
| s08 | Jk(a+<sup>weak</sup> b-) | Jk(a+b+) | *JK\*01W.06* | *JK\*02N.09<sup>†</sup>* |
| Novel null alleles |  |  |  |  |
| s09 | Jk(a+b-) | Jk(a+b+) | *JK\*01* | *JK\*02(c.725G>A)Null<sup>§†</sup>* |
| s10 | Jk(a-b+) | Jk(a+b+) | *JK\*01(c.119G>A)Null<sup>§</sup>* | *JK\*02<sup>†</sup>* |
| Novel structural variant |  |  |  |  |
| s02 | Jk(a-b+) | Jk(a+b+) | *JK\*01(Ex9_10del)Null<sup>§</sup>* | *JK\*02<sup>†</sup>* |
| s03 | Jk(a-b+) | Jk(a+b+) | *JK\*01(Ex9_10del)Null<sup>§</sup>* | *JK\*02<sup>†</sup>* |
| s07 | Jk(a-b+) | Jk(a+b+) | *JK\*01(Ex9_10del)Null<sup>§</sup>* | *JK\*02<sup>†</sup>* |

§ Novel alleles not yet described. † Allele harbors at least one additional silent SNV not reported in corresponding ISBT allele.

Three samples did not show any promising candidates when calling SNVs and short indels. However, SV calling identified a large 5,095 bp deletion in all three samples (Figure 2a). Using NG_011775.4 as reference, *cuteSV* suggested breakpoints located before position 64,898 bp in intron 8 and after position 69,992 bp in exon 10 (NG_011775.4:g.64898_69992del; NM_015865.7:c.947-1453_*2066del, Figure 2b). Bridge-PCR followed by Sanger sequencing confirmed exactness of these breakpoint positions (Figure 2d). In all three samples, the deletion was located on a *JK*01* allele (c.838G), and donors were *JK*01*/*JK*02* heterozygotes.

## Discussion

By employing a combination of LR-PCR and long-read nanopore sequencing, we successfully resolved all genotype-phenotype discrepancies in the Kidd blood group system observed over seven years of routine high-throughput donor genotyping. Our developed protocol allowed sequencing the complete coding region of the *SLC14A1* gene including introns as haplotypes, greatly facilitating the identification of the underlying genetic basis for the observed genotype-phenotype discrepancies and directly producing high-quality haplotype reference sequences for newly described *JK* alleles.

The majority of *JK* weak and null alleles listed in the ISBT tables are known to occur at very low frequencies across different populations [4,21,22], supporting the low number of discrepancies identified in our study. Specifically, nanopore sequencing allowed us to identify the following known SNVs: c.130G>A (rs2298720), c.191G>A (rs114362217), c.742G>A (rs763095261), c.871T>C (rs78242949) and c.956C>T (rs565898944). The SNVs could be unambiguously phased to the respective *JK*01/02* background allele, thus resulting in *JK*02W.03*, *JK*02N.09*, *JK*01W.05*, *JK*02N.06* and *JK*02N.08* alleles, respectively (Table 1). All these variants show minor allele frequencies (MAFs) in large-scale sequencing projects [23] that are in close agreement with the exceeding rarity observed in our study (MAFs < 0.001), with the exception of rs2298720, which is a common SNV (see discussion below). It is important to acknowledge that these variants (alleles) were only detected in *JK*01/02* heterozygotes of our donor population since discordant serology would not have appeared in homozygotes due to the masking effect of the wildtype allele.

Our workflow did not aim to detect variants that determine weak alleles, given that the transfusion recommendations for carriers of weak JK phenotypes are not different from those with normally expressed antigens. Consequently, weak agglutination levels were not considered as discrepant as long as in agreement with *JK01/02* genotyping. Over seven years of MALDI-TOF MS high-throughput donor genotyping, only two known weak alleles were flagged as suggestively causative for discrepant cases, i.e. resulting in the absence of observable levels of agglutination with respective anti-Jk antibodies. One was the aforementioned variant c.742G>A, defining the *JK*01W.05* allele. This allele is currently listed as ’weak’, but has already been reported to cause null phenotypes by others [24]. The other variant was c.130G>A, defining the *JK*02W.03* allele. This variant is a common SNV (MAF > 0.05; rs2298720), which also defines the most frequent weak allele on the *JK*01* background (*JK*01W.01*) along with several other weak alleles [13]. Since we did not observe further discrepancies pointing to weak alleles harboring this variant, there is little evidence that expression levels of *JK*02W.03* commonly fall below the detection threshold, pointing to an exceptional case reported here. Without phenotyping methods offering higher sensitivity, however, we cannot conclusively determine whether expression levels caused by these two presumably weak alleles were just below our detection threshold or if those two alleles indeed caused true null phenotypes in our samples. Efforts for in-depth phenotyping using absorption/elution techniques are ongoing.

Among the remaining cases that displayed genotype-phenotype discrepancies in our study, two were found with exonic SNVs likely defining new null alleles not yet listed by the ISBT [13]. One of these variants, c.725G>A (rs1197896884), is located in exon 7 and has been previously reported in two Europeans in the gnomAD [25] sequence collection of more than 125,000 sequenced exomes and whole genomes. The other variant, c.119G>A, is entirely novel. Notably, a singleton with a different base change (G>C) at the same position resulting in a different amino acid change (Gly40Ala) has been found in the gnomAD [25] sequence collection (rs1253002394), suggesting a tri-allelic SNV. Furthermore, it is noteworthy that a yet another change of the same amino acid (Gly40Ser, c.118G>A, rs145283450) is known to lead to the presence of null alleles *(JK*01N.17, JK*02N.21*) [13,26].

Our most remarkable finding in this study was the discovery of a large structural variant that was underlying three of the observed genotype-phenotype discrepancies. Our nanopore sequencing strategy revealed a large deletion of approximately 5 kb spanning from exon 9 to 10 on the *JK*01* allele. This deletion represents, to the best of our knowledge, the largest SV ever documented within the JK blood group system. Currently, the Kidd ISBT allele table only encompasses one deletion of ∼1.6 kb defining the *JK*01N.01* allele. First reported in Tunisian women [27], it has been sporadically reported in several other populations [18,28]. This deletion (c.1_341del) spans over exons 3 and 4 of the *SLC14A1* gene [27] and results in the lack of translation of the Kidd protein as the start codon is located in exon 3. Considering the complete absence of exon 9 and a portion of exon 10 in alleles carrying our newly discovered large deletion, it is highly probable that the resulting protein is also not functional. However, as in the case of the two aforementioned SNVs causing *JK*02W.03* and *JK*01W.05*, additional in-depth analyses using adsorption-elution or flow cytometric experiments [29] will be required to validate a true null phenotype of the three samples harboring the large deletion.

The apparent lack of described SVs in the Kidd system extends to almost all blood groups [13]. With the continuing accumulation of high-coverage whole genome sequencing datasets and improved algorithms for SV discovery, it has become clear that SVs appear to be more common in the human genome than previously thought. Depending on the population targeted and the sequencing technology used, between 7,500 and 22,600 SVs per genome were observed [30,31]. Additionally, SVs are responsible for an estimated ∼30% of rare heterozygous (MAF < 1%) gene inactivation events per individual [30]. Therefore, the extremely rare reports of SVs in blood group genes could simply be linked to the hitherto lack of adequate tools for their detection.

This study serves as a compelling example for illustrating the power of nanopore sequencing in resolving complex SVs that would have remained elusive using conventional sequencing methods. Indeed, using Sanger sequencing alone, the sequencing results for exon 9 and 10 of the three samples harbouring the deletion would have appeared as homozygous since only the wildtype allele would have been amplified and sequenced. In contrast, nanopore sequencing enabled us to precisely identify the breakpoints, which were subsequently verified using bridge-PCR.

With the continuing reduction in cost and the increase in accessible and easy-to-use bioinformatic tools, we forecast that long-read sequencing technologies will be more and more frequently used to overcome challenging diagnostic cases in blood group genetics.

### Author Contributions

Conceptualization, M.G., G.A.T., S.M. and M.P.M.-G.; methodology, M.G., G.A.T., N.T., L.S., S.S., N.L. and Y.M.; software, M.G. and G.A.T.; validation, M.G., G.A.T., N.T., L.S., S.M. and M.P.M.-G.; formal analysis, M.G. and G.A.T.; resources, C.E., B.M.F., C.G. and S.M.; writing – original draft preparation, M.G.; writing – review & editing, M.G., G.A.T. and M.P.M.-G.; visualization, M.G.; supervision, S.M. and M.P.M.-G. All authors read and approved the final manuscript.

## Funding Information

This research received no external funding.

### Institutional Review Board Statement

Not applicable. (According to the Swiss cantonal and national legislation, molecular blood group analyses are no subject to ethical authorization).

### Informed Consent Statement

General informed consent was obtained from all subjects involved in the study.

### Data Availability Statement

The identified 5,095 bp deletion (NG_011775.4:g.64898_69992del; NM_015865.7:c.947-1453_*2066del) on the *JK*01* haplotype has been submitted to ClinVar (accession number: VCV001202625.1). Full-length haplotype sequences of all sequenced alleles, covering the entire coding region of SLC14A1, have been submitted to GenBank (accession numbers: PP034563-PP034582).

